# A High Sensitivity Assay of UBE3A Ubiquitin Ligase Activity

**DOI:** 10.1101/2025.01.05.631408

**Authors:** Linna Han, Z. Begum Yagci, Albert J. Keung

**Affiliations:** Department of Chemical and Biomolecular Engineering, North Carolina State University, Campus Box 7905, Raleigh, NC 27695-7905, U.S.A

**Keywords:** Biochemical assay, neurodevelopmental disorder, therapeutic endpoint, biomarker, Angelman Syndrome, Autism Spectrum Disorder, Dup15q Syndrome

## Abstract

UBE3A is an E3 ubiquitin ligase associated with several neurodevelopmental disorders. The development of several preclinical therapeutic approaches involving UBE3A, such as gene therapy, enzyme replacement therapy, and epigenetic reactivation, require the detection of its ubiquitin ligase activity. Prior commercial assays leveraged Western Blotting to detect shifts in substrate size due to ubiquitination, but these suffered from long assay times and have also been discontinued. Here we develop a new assay that quantifies UBE3A activity. It measures the fluorescence intensity of ubiquitinated p53 substrates with a microplate reader, eliminating the need for Western Blot antibodies and instruments, and enables detection in just 1 hour. The assay is fast, cost-effective, low noise, and uses components with long shelf lives. Importantly, it is also highly sensitive, detecting UBE3A levels as low as 1nM, similar to that observed in human and mouse cerebrospinal fluid. It also differentiates the activity of wild-type UBE3A and catalytic mutants. We also design a p53 substrate with a triple-epitope tag HIS-HA-CMYC on the N terminus, which allows for versatile detection of UBE3A activity from diverse natural and engineered sources. This new assay provides a timely solution for growing needs in preclinical validation, quality control, endpoint measurements for clinical trials, and downstream manufacturing testing and validation.

## Introduction

Ubiquitination is a post-translational modification with diverse regulatory functions ranging from protein degradation and gene expression to subcellular localization and kinase activation [1,2,3,4]. Three key enzymes are involved in the ubiquitination cascade pathway: ubiquitin-activating enzyme (E1), ubiquitin-conjugating enzyme (E2), and ubiquitin ligase (E3) [5]. E3 ubiquitin ligases, in particular, modulate various biological and developmental processes [6].

The *UBE3A* gene encodes a specific 100 kDa E3 ubiquitin ligase belonging to the HECT (homologous to E6-AP C-terminus) domain family. *UBE3A* is expressed from both the paternal and maternal alleles of chromosome 15 in most tissues. However, in neurons, paternal *UBE3A* is imprinted and silenced [7]. Deletion or loss of activity of the maternally inherited *UBE3A* results in an absence or reduction of UBE3A protein in neural cell types [8, 9, 10, 11, 12, 13] and is linked to autism spectrum disorders (ASDs) and Angelman syndrome (AS) [14, 15]. Its ubiquitin ligase activity has been found to be particularly important [16, 17, 18, 19, 20]. These findings have motivated several preclinical therapeutic approaches currently being developed for Angelman Syndrome including enzyme replacement therapies (ERT), gene replacement, and epigenetic reactivation [21, 22, 23, 24, 25, 26, 27, 28, 29].

Given the biological and therapeutic importance of UBE3A, there is a need for tools to detect the ubiquitin ligase activity of UBE3A for mechanistic studies, biomarker measurements, as well as for manufacturing and clinical endpoints measurements. Existing assays include Western Blot analysis of size shifts caused by ubiquitination of a substrate [30, 31, 32, 33] or on intracellular luciferase and WNT activity assays [34, 35] performed in HEK cells. Our goal was to develop a highly controlled in vitro assay that avoids the use of cell-based measurements and Western Blotting, and that is fast, sensitive, and cost-effective. The resultant assay possesses high sensitivity that is capable of detecting UBE3A as low as 1 nM. This is at the expected levels of UBE3A found in cerebrospinal fluid where it is expected to be at lower levels than in cells and tissue but is accessible for biomarker and clinical sampling. The assay can also be completed in one hour with a simple plate reader and standard molecular biology equipment. As described here, this assay does not detect specific activity towards substrates other than p53, and it does not account for substrate specificities conferred by different combinations of E1 and E2 proteins. Furthermore, in complex samples, it remains unclear if it can distinguish UBE3A activity from the activity of other E3 ligases that may be present. Its utility for mechanistic work is therefore currently limited, with its advantages supporting primarily therapeutic development. However, modifications to the assay could direct it towards probing more specific activities of UBE3A in the future.

## Results

### A plate-reader based assay detects ubiquitin ligase activity of UBE3A

A central aim of this work was to replace the use of Western Blot analysis with commonly used plate readers. Towards this goal, we developed an initial assay where fluorescein conjugated ubiquitin is combined with an epitope-tagged substrate, UBE1, UBE2, Mg^2+^, ATP, and an experimental sample containing commercial GST-tagged UBE3A or no UBE3A (Table 1). If UBE3A is present, it will ligate the fluorescein-ubiquitin to the substrate. Subsequently, a magnetic bead coated with an antibody against the epitope tag is introduced to pull down the substrate and separate it from free fluorescein-ubiquitin (Figure 1a). ATP is included, as E1 enzymes activate ubiquitin in an ATP dependent mechanism. p53 is used as the substrate [36] and is HIS-tagged to facilitate magnetic pulldown. The purity of all protein components purity was checked by a SDS-PAGE gel (Supplemental Fig. S1).

**Table 1.** Assay Components. Initial conditions used in Figures 2-4 unless otherwise noted. Assay components are listed in the order they were added to the reaction mixture. The concentration of UBE3A varies according to the experiment performed and are noted in figure captions, but the concentrations of the other components remain the same. 1 μM HPV-E6 was also included in experiments in Figures 2-4 and was added directly after UBE3A was added.

|  | Working Concentration or Amount |
| --- | --- |
| dH<sub>2</sub>O |  |
| 10*Rx Buffer | 50 mM HEPES, pH 8.0, 50mM NaCl |
| Mg<sup>2+</sup> | 10 mM |
| ATP | 10 mM |
| UBE1 (UBA1) | 100 nM |
| UBE2 (UB2L3) | 1 μM |
| UBE3A | 200 nM |
| HIS-FLAG-p53 | 5 μg |
| Fluorescein-ubiquitin | 2.5 μM |

**Figure 1.**
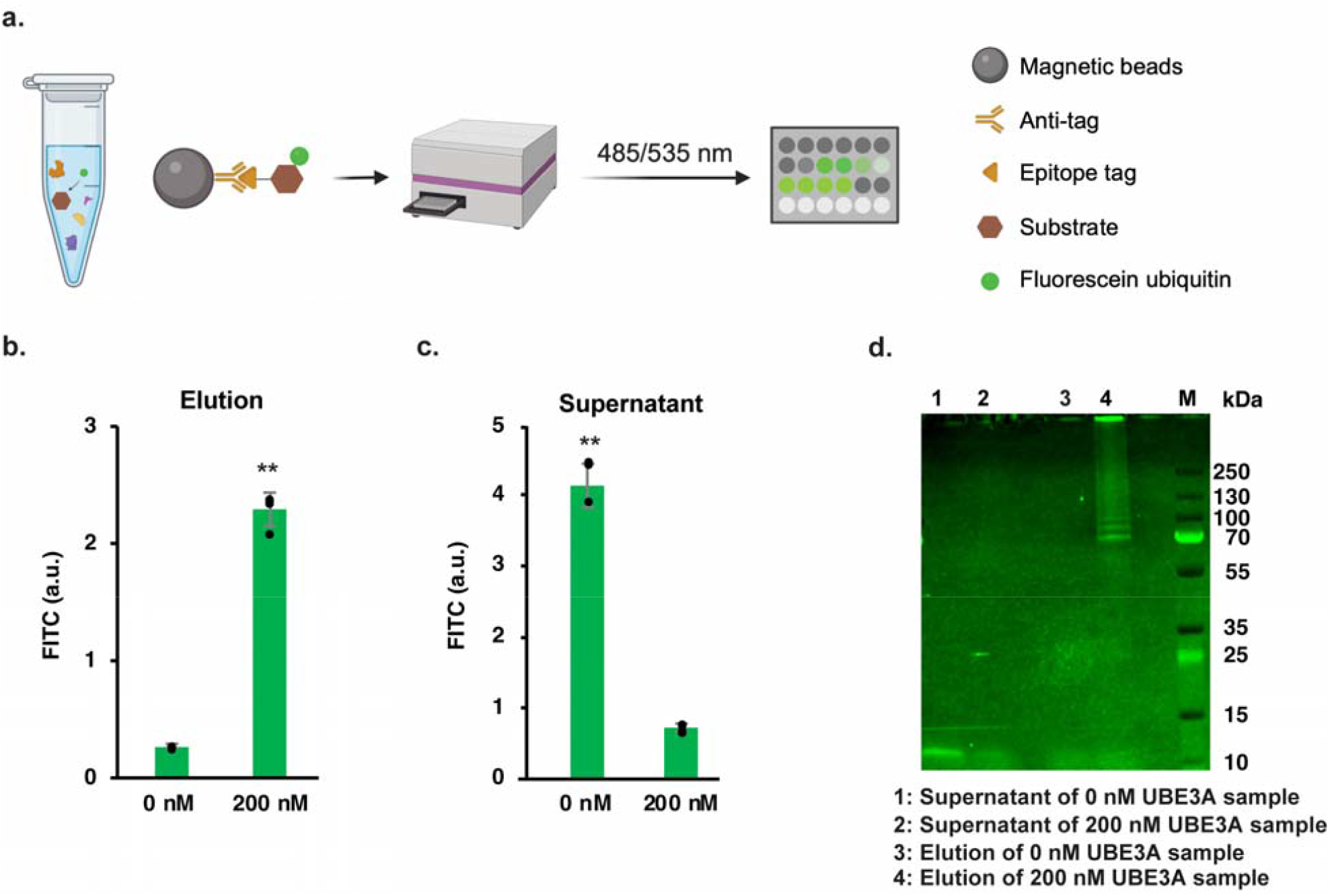
A plate-reader based assay detects ubiquitin ligase activity of UBE3A. (a) Schematic of assay (Created with BioRender.com). (b,c) Plate reader detection of FITC fluorescence for the elution (b) and supernatant (c) from the assay using 200 nM UBE3A for a 60 min incubation time. (d) SDS-PAGE visualization of fluorescein-ubiquitin conjugated to p53 appearing only in the elution of the sample with UBE3A added. N=4 separate replicate reactions. Error bars represent standard deviation. **p<0.01 relative to sample without UBE3A. ANOVA with Tukey-Kramer post-hoc.

To test this assay, samples with or without 200 nM UBE3A were incubated in the reaction mixture at 37 °C for 60 minutes followed by addition of anti-HIS magnetic beads to pull out p53. The fluorescence intensity of the supernatant, as well as the subsequent elution of the p53 off the magnetic bead, were quantified by a fluorescence plate reader (Fig. 1b). As expected, when UBE3A was present, most of the fluorescein-ubiquitin was ligated to p53 and appeared in the elution while there was little free fluorescein-ubiquitin remaining in the supernatant. Both the reaction and elution buffers exhibited minimal background fluorescence. We also verified the pull-down assay with an SDS-PAGE analysis visualizing the fluorescein-ubiquitinated p53 only in elution of the sample containing UBE3A (Fig. 1c).

### Inclusion of HPV-E6 and reducing reaction volumes improves assay efficiency

Biochemical assays are often expensive due to the use of recombinant proteins and specialty chemicals. We therefore sought to increase the efficiency of the assay through three modifications. First, we asked if HPV-E6 could increase the sensitivity of the assay. E6 (Early protein 6) is a viral protein produced in cells infected with the Human Papillomavirus. E6 forms a complex with the host cell UBE3A generating a ligase activity that polyubiquitinates target substrates [37]. When included in the reaction, HPV-E6 did indeed boost the assay signal by nearly fivefold compared to a sample with the same concentration of UBE3A but lacking HPV-E6 (Fig. 2a). It was reported that UBE3A can ubiquitinate itself, a process influenced by the presence of p53, which forms protein complexes with UBE3A and HPV-E6 [38]. To evaluate the specificity of this assay, we performed a ubiquitination reaction and divided the reaction mixture into two parts, using anti-HIS and anti-GST magnetic beads to separately pull down HIS-tagged p53 and GST-tagged UBE3A. While GST tagged UBE3A exhibited a slight fluorescence signal, it did not differ significantly from the control sample without UBE3A (Supplemental Fig. S2). Second, we then asked if the volume of the reaction could be scaled down to preserve reagents. We found that the reaction could be scaled down from 50 μL to 25 μL or 10 μL while preserving the sensitivity of the assay (Fig. 2b). The enzyme concentration was maintained across all three volume conditions, meaning less total enzyme was required by scaling the volume down. The substrate, p53, was maintained at a constant 5ug total amount across all three volumes. Finally, we asked if UBE3A ubiquitinates FADD (FAS-associated death domain protein), which like p53 is a binding partner of E6 [39, 40]. We compared its level of ubiquitination to p53 but found it was substantially weaker and not statistically significant compared to a sample with no UBE3A added (Fig. 2c). This provided some evidence for the specificity of the assay for UBE3A activity. Consequently, we employed p53 as the substrate for all subsequent assays.

**Figure 2.**
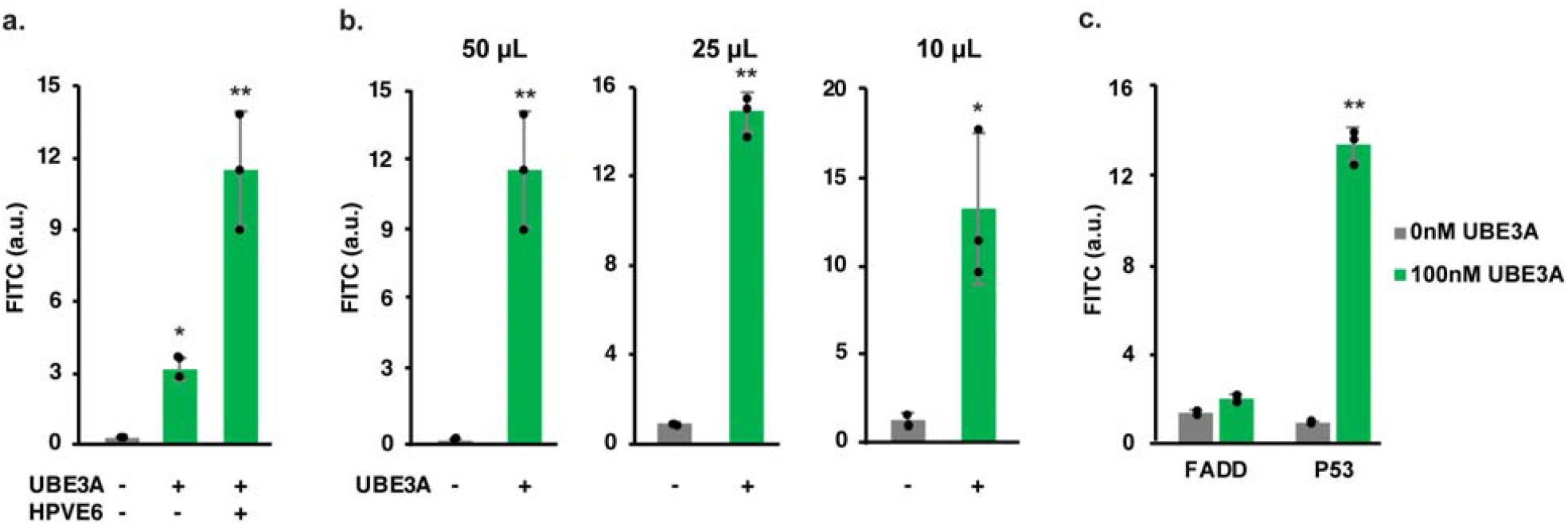
Inclusion of HPV-E6 and reducing reaction volumes improves assay efficiency. (a) 1 μM HPV-E6 boosts ubiquitination by almost five folds. (b) Reduction in reaction volume maintains assay sensitivity. (c) FADD is not a substrate of UBE3A. (a-c) UBE3A levels were 200 nM and incubation times were 60 min for all experiments in this figure. N=3 separate replicate reactions. Error bars represent standard deviation. **p<0.01, *p<0.05 relative to sample without UBE3A. ANOVA with Tukey-Kramer post-hoc in panel (a). Two-tailed t-test comparison to sample without UBE3A in panels (b) and (c). (b and c) 1 μM HPV-E6 was included in all of these reactions.

### p53 with multiple epitope tags provides assay versatility

In the assay, we used commercially available HIS-tagged p53, and the HIS tag to pull out the substrate after the ubiquitination reaction. However, UBE3A protein generated by different therapeutic modalities including for enzyme replacement therapy or gene therapy, may include epitope tags as well which could cross react with an identical epitope used for pulldown of the substrate. We therefore designed a custom p53 substrate with three epitope tags, HIS-HA-CMYC, appended to the N-terminus of p53. This version of substrate could be universally used by researchers by choosing antibodies and magnetic beads that bind an epitope present on p53 but not on any engineered UBE3A of interest. We tested that this substrate maintains activity in the assay when using anti-HIS, anti-HA, or anti-CMYC magnetic beads to pull out p53 following a ubiquitination reaction. All three epitope tags worked well (Fig. 3). The anti-CMYC assay yielded slightly better signal but costs three times that of anti-HIS beads. We therefore used anti-HIS beads for most of the remaining work except in cases where UBE3A contained a HIS tag.

**Figure 3.**
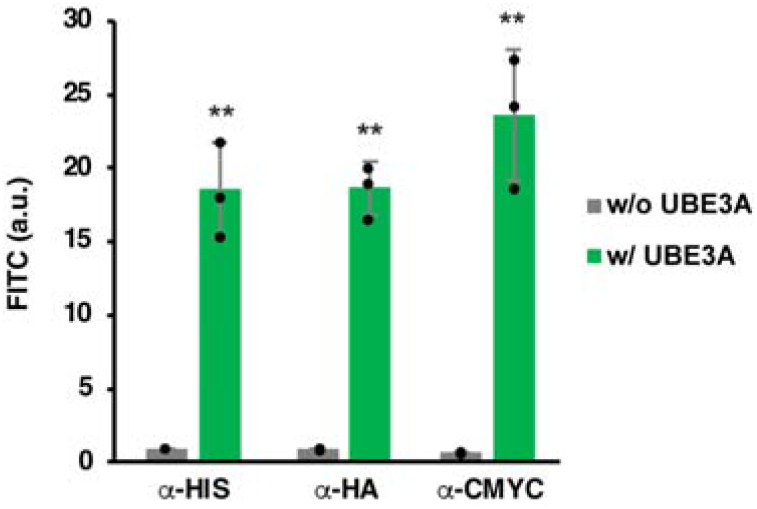
p53 with multiple epitope tags provides assay versatility. (a) SDS-PAGE gel indicating successful production of a p53 substrate with triple HIS-HA-CMYC epitope tags. (b) The assay performs similarly when using magnetic beads pulling down the p53 substrate using each of the three epitope tags. N=3 separate replicate reactions. Error bars represent standard deviation. **p<0.01, Two-tailed t-test relative to sample without UBE3A.

### Optimization of assay conditions for different target UBE3A concentrations

Users may have different applications ranging from testing recombinantly produced UBE3A to assaying UBE3A activity within biological samples. These applications could have substantially different demands on assay sensitivity. We therefore first profiled the assay at four different reaction incubation times (5, 30, 60, 90 minutes) and at four concentrations of UBE3A (1, 10, 25, 50 nM). The reaction was stopped by putting reaction tubes on the ice after the ubiquitination reaction and incubating with the magnetic beads at 4 °C. The signal from 1, 10 and 25 nM UBE3A increased with extended incubation time, while 50nM UBE3A saturated at 60 minutes (Fig. 4a).

**Figure 4.**
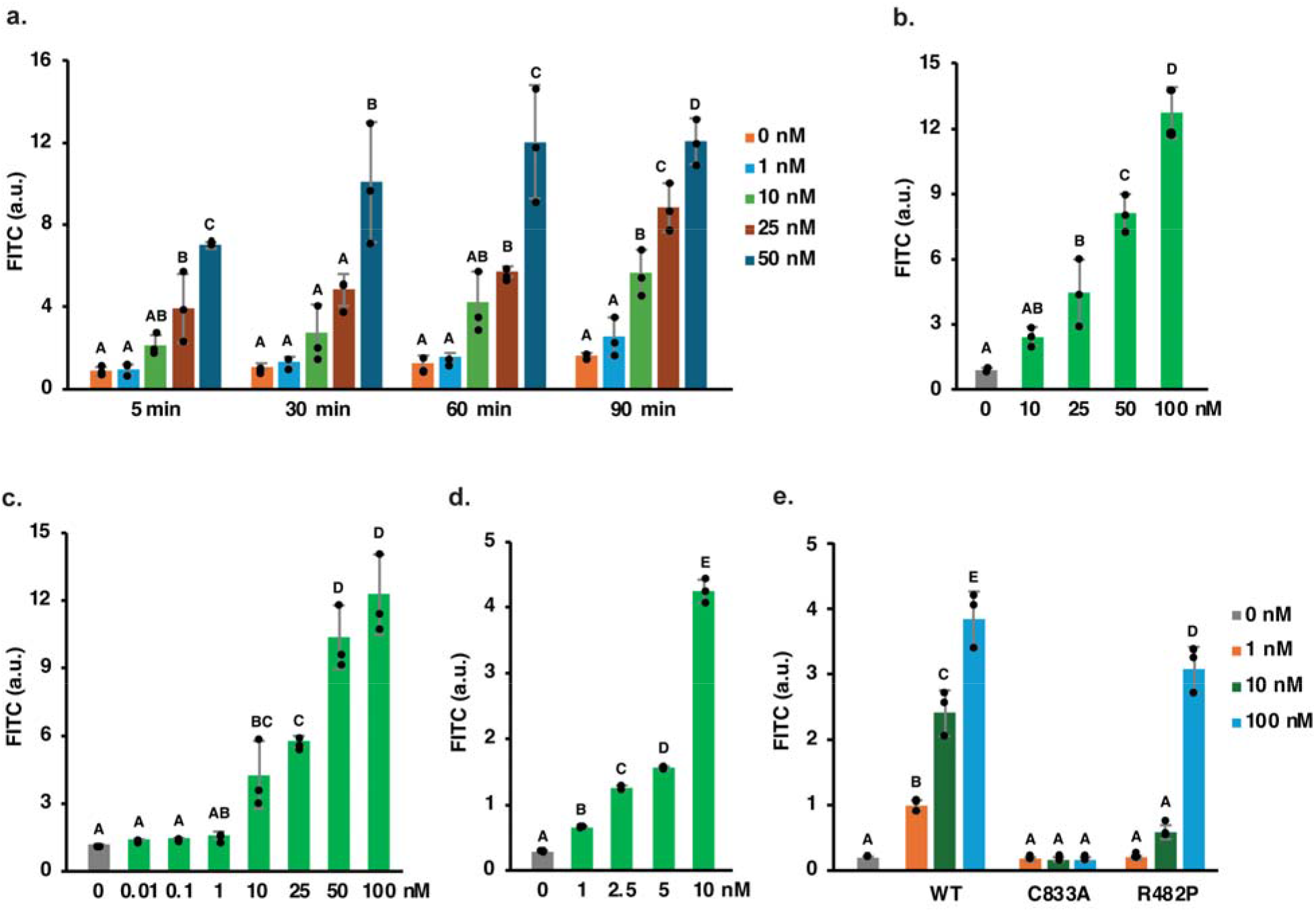
Optimization of assay conditions for different target UBE3A concentrations. (a) Fluorescence signal from the assay performed with four incubation times and five concentrations of UBE3A. Fluorescence signal from the assay performed with (b) 5 minute and (c) 60 minute incubation times, with incubation of the magnetic beads at 4 °C for 30 minutes. (d) Fluorescence signal from the assay performed with 10 μM fluorescein-ubiquitin over a 60 minute incubation time. (e) The assay detects lower ubiquitination by the R482P UBE3A mutant and no ligase activity by the C833A UBE3A, with assay conditions of 1, 10 and 100nM UBE3A. In-house UBE3A variants (WT, R482P and C833A) were used in Figure 4e, along with anti-CMYC magnetic beads for the pulldown assay. N=3 separate replicate reactions. Error bars represent standard deviation. Samples with the same letters are not statistically significant from each other. Otherwise, p<0.05 relative to samples without the same letter. ANOVA with Tukey-Kramer post-hoc.

In some applications like assaying recombinant UBE3A protein, speed would be preferable and there would not be a substantial limit on the amount of UBE3A available for testing. We tested what concentrations could be detected by the assay with only a 5 minute reaction incubation and found 25 nM UBE3A and higher could be detected as statistically different from the negative control (Fig. 4b). Including the other steps, the entire assay could then be completed in 1 hour. In other applications, the amount of UBE3A protein may be low. We therefore tested a broader range concentrations of UBE3A, finding that a 60-minute incubation was only able to distinguish 10 nM UBE3A and higher (Fig. 4c).

We further investigated the effect of fluorescein-ubiquitin on UBE3A detection sensitivity. A dose-response curve was generated using 1, 5, 10, 50, and 100 μM fluorescein-ubiquitin with 1 and 10 nM UBE3A separately, while 0 nM UBE3A served as a negative control. The highest fluorescence intensity peak for both concentrations of UBE3A occurred at 10 μM fluorescein-ubiquitin, with higher concentrations leading to a decrease in FITC signal (Fig. S3). Subsequently we set up assays with 10 μM fluorescein-ubiquitin to measure UBE3A activity at 1, 2.5, 5 and 10 nM over a 60-minute incubation.

Compared to 90 minute assay results with 2.5 μM fluorescein-ubiquitin, this optimized fluorescein-ubiquitin concentration significantly boosted UBE3A signal, enabling the detection and resolution of 1, 2.5, 5, and 10 nM UBE3A (Fig. 4e).

Lastly, we tested whether the assay could distinguish wild type UBE3A with a catalytic mutant R482P [35] and the ligase dead mutant C833A [41] through dose-response assays. All three proteins were recombinantly produced and purified. They were all tagged with HIS; therefore, p53 was pulled down by anti-CMYC beads to avoid cross-interaction. Using a 60 minute incubation with 1, 10, 100nM UBE3A separately, the assay successfully differentiated wildtype UBE3A from the R482P mutant, particularly at 1 and 10nM concentrations. C833A displayed no ligase activity at any of the three concentrations (Fig. 4f).

### Application of the assay on the cell lysates

To test the assay with biological samples, we performed an experiment in which we collected cell lysates from H9_*WT*_ and H9_*UBE3Am-/p-*_ human stem cells. In our previous study, we confirmed the absence of *UBE3A* in H9_*UBE3Am-/p-*_ (generated by Chamberlain group) [10, 13, 42, 43, 44]. There are many commercially available cell lysis buffers. We chose Pierce™ IP lysis buffer and M-PER™ mammalian protein extraction reagent to test their compatibility with our assay. IP lysis buffer showed better fluorescence signal in our assay compared to M-PER reagent (Fig. S4a) indicating that it might work better with the assay. In addition, there are situations where concentration of dilute samples would help increase signal. Hence, we used Amicon® ultra centrifugal filters. As expected, an increased signal was detected when the sample was concentrated (Fig. S4b). Hence, we decided to use IP lysis buffer and ultra centrifugal filters for our next experiment in which we used 3 biological replicates for each cell line. Our results showed that the signal from H9_*WT*_ cell lysate was significantly higher than that from H9_*UBE3Am-/p-*_(Fig. 5a), with the background signal without any sample added was ∼ 5% of the difference in signal between the H9_*WT*_ and H9_*UBE3Am-/p-*_ cell lysates. Taken together, these results show the potential of our assay to be used with biological samples. This experiment with human stem cell lysates showed that IP lysis buffer and ultra centrifugal filters could work with this assay. However, M-PER and other buffers should not be eliminated as an option since it might work better with different types of samples.

**Figure 5.**
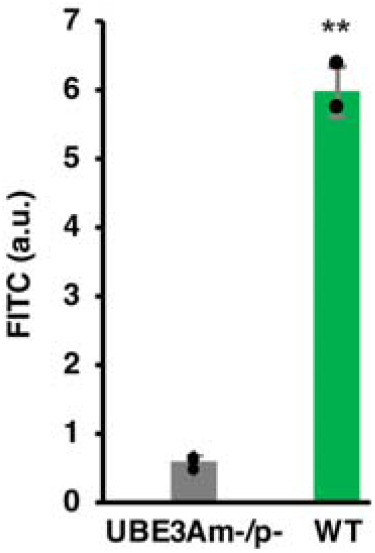
UBE3A activity is detected in human stem cell lysates. Fluorescence signal from the assay performed with the cell lysates obtained from H9_*WT*_ and H9_*UBE3Am-/p-*_ human stem cells. The fluorescence intensity signal of the negative control (lysis buffer only) was first subtracted from the cell lysate readings. The readings were then equilibrated to the same total protein amount loaded into the assay. The negative control contains filter-concentrated IP lysis buffer along with other assay components except for UBE3A. N=3 separate biological samples collected from different wells of a 6-well plate. Error bars represent standard deviation. **p<0.01 relative to UBE3A-double knocked out sample. Two-tailed t-test was performed to calculate p-value.

### *In vitro* ubiquitin ligase assay protocol

**Step 1:** In a low-binding microcentrifuge tube, combine each component listed **in the order** shown in Table 2. For a negative control reaction, replace the UBE3A with dH2O or with your specific negative control. The total reaction volume for each sample can start as low as 10uL and can be scaled up accordingly.

**Table 2.** Optimized ubiquitin conjugation assay components. Optimized and recommended ubiquitin conjugation assay components and working concentrations.

| Components | Reagents | Working Concentration |
| --- | --- | --- |
| Buffer | 50 mM HEPES, pH 7.5, 50 mM NaCl | 50 mM |
| Energy | Mg<sup>2+</sup>-ATP | 10 mM |
| Activating enzyme | UBE1 (UBA1) | 100 nM |
| Conjugating enzyme | UBE2L3 | 1 µM |
| Ligase | UBE3A/E6AP | Sample |
| Ligase complex component | HPV 16-E6 | 1 µM |
| Substrate | HIS-HA-CMYC-p53 | 5 µg |
| Ubiquitin | Fluorescein-ubiquitin | 10 µM |

\*Once you add the fluorescein-ubiquitin, the reaction begins.

**Step 2:** Briefly centrifuge the mixture at 2000g, incubate in a 37°C water bath for 60 minutes (the time can be shortened to as low as 5 minutes or extended to 90 minutes, depending on your objective). For situations where you are testing the effect of incubation time, the reaction can be terminated by placing the reaction samples on ice. **Otherwise, keep all steps at room temperature**.

**Step 3:** Incubate reaction mixtures with pre-washed HisPur Ni-NTA magnetic beads (Fisher 88831) or anti-CMYC (Fisher 88842) or anti-HA (Fisher 88836) beads in 400 μL Equilibration Buffer (25 mM HEPES, 0.15M NaCl, 0.25% Tween-20 Detergent)** on an end-over-end rotator for 30 min.

**Step 4: Briefly centrifuge to remove potential fluorescein-ubiquitin on the lid**. Collect the beads by placing the reaction tube on a magnetic stand, save the supernatants for subsequent fluorescence intensity analysis.

**Step 5:** Wash the beads by adding Wash Buffer (25 mM HEPES, 0.15M NaCl, 0.25% Tween-20 Detergent, 20 mM Imidazole)** to the tube, then vortex for 10 seconds to mix. Collect the beads by placing the tube on a magnetic stand, then remove and discard the supernatant. Repeat Step 5 once.

**Step 6:** Add 50uL Elution Buffer (25 mM HEPES, 0.15M NaCl, 0.25% Tween-20 Detergent, 200 mM Imidazole)** to each tube, vortex for 15 seconds. Incubate the beads at room temperature for 15 min with vortexing the tube every 5 minutes.

**Step 7:** Magnetically separate the beads and save the supernatant that now contains the eluted ubiquitinated-p53 to a **384-well black plate** (Corning 3575, or other black plate). Measure fluorescence intensity at ex485/em535nm on a microplate reader.

**Anti-HA and anti-CMYC magnetic beads use 1 x TBS-T (25 mM Tris, 0.15M NaCl, 0.25% Tween-20 Detergent) as the Equilibration Buffer, 5 x TBS-T (125 mM Tris, 0.75M NaCl, 1.25% Tween-20 Detergent) as the Wash Buffer, and 50 mM NaOH as the Elution Buffer.

To prepare UBE3A-containing cell lysate:

**Step 0-1:** Follow the manufacturers’ protocols for Pierce™ IP lysis buffer (Fisher Scientific PI87787) or another cell lysis buffer that does **NOT** contain denaturing components. Begin by washing the cells with ice cold Gibco™ DPBS (Fisher Scientific 14190250). Add 300-400µl IP lysis buffer to each well and gently shake the cells on ice for 20-30minutes. Then, scrape cells and centrifuge the cell solutions at 13,000g for 10min at 4°C. Collect the supernatant and store it at -80°C (long-term storage) or at 4°C (short-term storage) until further analysis.

**Step 0-2:** Follow the manufacturer’s instructions for the Amicon® ultra centrifugal filters (MilliporeSigma, UFC505096, 50kDa MWCO, 0.5ml) to concentrate the cell lysates to the desired volume. For example, to prepare a 25uL reaction mixture for Step 1, concentrate the cell lysates to approximately 15uL. Assume that the protein loss is equivalent across samples prepared at the same time.

## Discussion

In this study, we developed a new UBE3A detection assay that is quantitative, versatile to diverse sensitivity ranges and engineered UBE3A proteins, and rapid and facile to perform. It uses widely available microplate readers to detect fluorescence signals. It aims to replace Western Blot assays that require more equipment and multiple lengthy processing steps. This assay is also fast and cost-effective (Table 3).

**Table 3.**
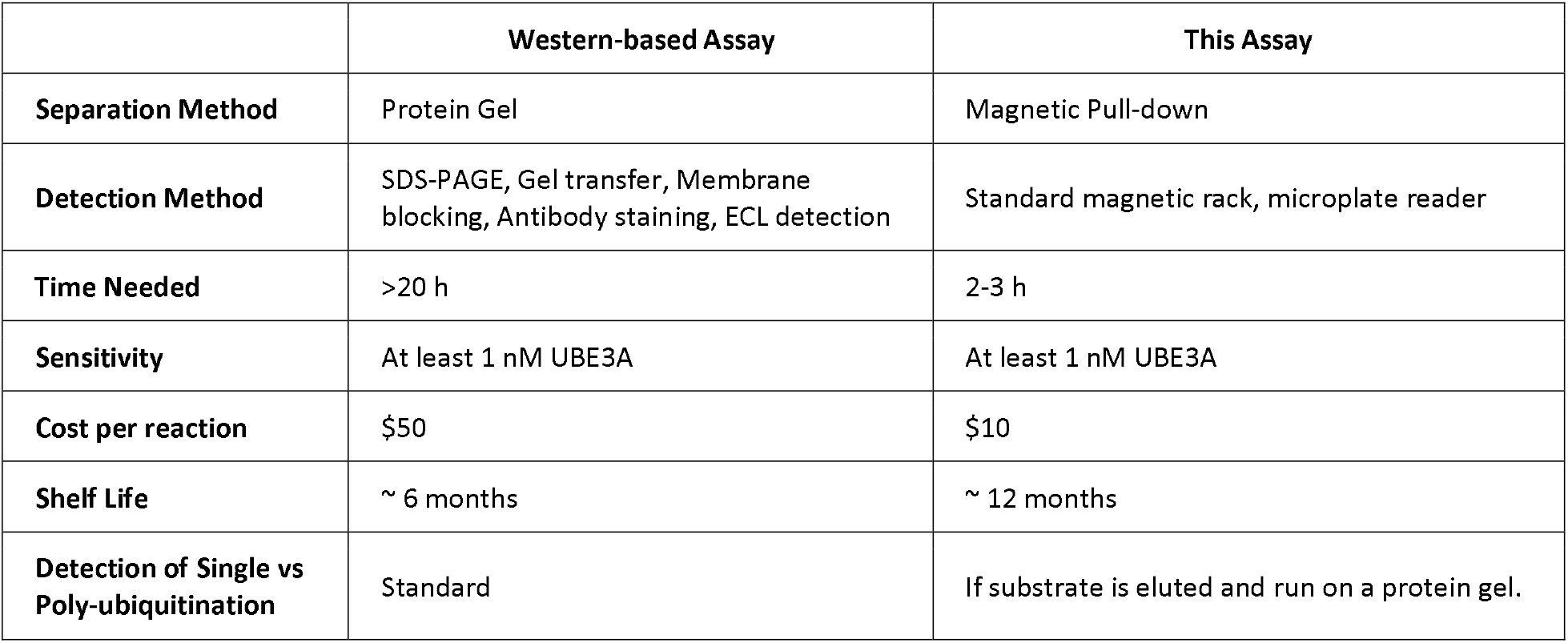
Technical Differentiators. Technical differentiators of this assay compared to western blot assays.

|  | Western-based Assay | This Assay |
| --- | --- | --- |
| **Separation Method** | Protein Gel | Magnetic Pull-down |
| **Detection Method** | SDS-PAGE, Gel transfer, Membrane blocking, Antibody staining, ECL detection | Standard magnetic rack, microplate reader |
| **Time Needed** | >20 h | 2-3 h |
| **Sensitivity** | At least 1 nM UBE3A | At least 1 nM UBE3A |
| **Cost per reaction** | \$50 | \$10 |
| **Shelf Life** | ~ 6 months | ~ 12 months |
| **Detection of Single vs Poly-ubiquitination** | Standard | If substrate is eluted and run on a proteingel. |

Another highlight of this assay is its exceptional sensitivity. Following optimization, it can detect UBE3A levels as low as 1 nM, the concentration expected in human cerebrospinal fluid CSF. Dodge and colleagues estimated the amount of UBE3A protein in human CSF to be ∼100 ± 7 ng/mL [45], which corresponds approximately to 1 nM. CSF sampling is of potential importance, as it represents the only widely applicable methodology for obtaining information on free drug concentrations within the individual human brain [46] and could therefore be used for assaying therapeutic responses and biomarker levels in clinical studies related to dysregulated UBE3A including for Angelman Syndrome, Dup15Q, and Autism Spectrum Disorder.

There were several parameters that most affected the performance, sensitivity, and robustness of the assay, and that could be further tuned to fit specific needs researchers may have. The sample amount as well as the amount of p53 and cofactor enzymes directly impacted the sensitivity and signal of the assay, albeit with a tradeoff in cost of cofactors, and potential limitations in sample amount available. The incubation time up to 30-60 minutes helped improve signal. Also, being consistent with the temperature of samples is important if shorter incubation times are used as variations in the time to equilibrate temperatures for different samples could introduce noise and bias. We also noticed that the age of UBE3A protein (and hence future experimental samples) did seem to affect the sensitivity of the assay, even when stored at -80 °C, and therefore is a consideration when performing longitudinal studies collecting samples from distantly separated time points.

The versatility of the system across low and high concentration ranges, and the ability to work with various engineered UBE3A proteins also provides potential utility for research in cell biology, neuropharmacology, and neuroscience. However, as described here, this assay does not detect specific activity towards substrates other than p53 and does not account for substrate specificities conferred by different combinations of E1 and E2 proteins. Western blot, WNT activity, and direct in vitro substrate ubiquitination assays (typically detected via Western blot) have been previously used to detect UBE3A activity and substrate specific ubiquitination. These methods have the advantages of being able to also directly detect changes in ubiquitination of specific substrates in the cellular context as well as in vitro, and also detect polyubiquitination of substrates, but suffer from lengthy procedures and pleiotropic effects that cannot always be decoupled from other ubiquitin ligases or signaling crosstalk. Some of these methods and the method described in this work could be adjusted towards specific E1 and E2 combinations or towards specific substrates of interest. Ultimately the ability of methods that detect activities directly within cells provides advantages for mechanistic studies, while the method described herein may be best suited for rapid and robust assessment of general UBE3A activity for purposes of downstream therapeutic development.

Future work could broaden the scope of this assay to detect all specific E1 and E2 combinations, as well as all known UBE3A substrates. It will be a challenging combinatorial problem; however, if solved, it could yield simultaneously interesting biological knowledge while providing a more comprehensive and complete picture of UBE3A activity. It is not clear whether a therapeutic enzyme replacement would exhibit activity towards one substrate but not another, given the proper E1 and E2 context. Presumably disruption of ligase activity during protein production or purification would generally impact activity for all substrates, but it is possible other portions of the enzyme conferring interaction specificities could be selectively disrupted.

## Materials and Methods

### Cloning, expression and purification of recombinant proteins

The E.coli codon-optimized gene coding for HIS-HA-CMYC-p53 was ordered from Twist and subcloned into pET (Addgene 29711) via ligation. Two gene fragments of wild type UBE3A isoform I were assembled into the pET vector with N-terminal Sumo and 6xHIS tags. UBE3A(R482P) and UBE3A (C833A) were created using site-directed mutagenesis. Proteins were produced in BL21(DE3) cells and purified by FPLC IMAC cartridges. The eluted proteins were concentrated using Amicon Ultra filters (30 kDa and 100 kDa MW cutoff separately) and stored at -80°C. Proteins were analyzed via SDS-PAGE (Supplemental Figure S1).

### Ubiquitin conjugation and pull-down assay

In low-bind microcentrifuge tubes, each component was added in the indicated order and concentration listed in Table 1. 1 μM HPV-E6 was added after UBE3A. UBE1 (UBA1, E-305-025), UB2L3 (E2-640-100), HPV-E6 (AP-120-025), fluorescein-ubiquitin (U-580-050) and HIS-FLAG-p53 (SP-452-020) are from R&D. Commercial GST-UBE3A (H00007337P01) is from Abnova. ATP solution (100mM) (FERR0441) is from Fisher. Negative control reactions were prepared by replacing UBE3A with dH2O. After a brief centrifugation at 2000g, the mixture was incubated in a 37 °C water bath for 5 minutes to 2 hours. The reaction was terminated by placing the reaction samples on ice unless otherwise described. Reaction mixtures were then incubated with pre-washed HisPur Ni-NTA magnetic beads (Fisher 88831) or anti-CMYC (Fisher 88842) or anti-HA (Fisher 88836) beads in Equilibration Buffer (25 mM HEPES, 0.15M NaCl, 0.25% Tween-20 Detergent) at 4 °C for 30 min with mixing, except where specified in the text at room temperature for 60 min. Supernatants were removed and saved for subsequent fluorescence detection. The beads were washed twice with Wash Buffer (25 mM HEPES, 0.15M NaCl, 0.25% Tween-20 Detergent, 20 mM Imidazole), and ubiquitinated p53 was eluted off the beads with Elution Buffer (25 mM HEPES, 0.15M NaCl, 0.25% Tween-20 Detergent, 200 mM Imidazole). The fluorescence intensity of the supernatants and elution were quantified on a TECAN plate reader. Anti-HA and anti-CMYC magnetic beads used 1 x TBS-T (25 mM Tris, 0.15M NaCl, 0.25% Tween-20 Detergent) as the Equilibration Buffer, 5 x TBS-T (125 mM Tris, 0.75M NaCl, 1.25% Tween-20 Detergent) as the Wash Buffer, and 50 mM NaOH as the Elution Buffer.

When checking for the possibility of UBE3A self-ubiquitination, the reaction mixture was separated into two parts, using anti-HIS and anti-GST magnetic beads to separately pull down HIS-tagged p53 and GST-tagged UBE3A. To pulldown GST-tagged UBE3A, the reaction mixture was incubated and washed by Equilibration/Wash Buffer (125 mM Tris-HCl, 150 mM NaCl, 1 mM DTT, 1 mM EDTA, pH 7.4). The commercial GST-UBE3A was eluted off the anti-GST magnetic beads (Fisher, 78601) with Elution Buffer (50 mM reduced glutathione) (Fisher, 78259).

For the assay performed on the cell lysates, the protocol described above was used. 25µl was chosen as the reaction volume. The reaction components used along with their approximate working concentrations were as follows: 10x reaction buffer (10 mM), Mg2+/ATP (10 mM), UBE1 (100nM), UBE2 (1 µM), HPV-E6 (1.5 µM), p53 (7.5 µg), ubiquitin-fluorescein (10 µM). The volume of the lysate sample and buffers used for the negative control was 12.75 µl. Mg2+/ATP (Enzo Life Sciences BMLEW98050100) was used instead of adding ATP and Mg^2+^, separately. This was done to allow more sample volume to be added. Negative control reactions were prepared by replacing UBE3A with IP lysis buffer, M-PER reagent or protein storage buffer (500mM HEPES, 150mM KCl, 10% Glycerol). For Fig 5, the IP lysis buffer was filter-concentrated before adding it to the control reaction mixture. The samples were incubated in a 37°C water bath for an hour. HisPur Ni-NTA magnetic beads (Fisher 88831) were used.

### SDS-PAGE

Proteins were mixed with reducing reagent and sample loading buffer. After incubation at 95 °C for 5 minutes, the samples were loaded into a Bis-Tris gel for SDS-PAGE analysis. Gel staining followed the Imperial Protein Stain protocol (Fisher PI24615). The elution and supernatant samples were mixed with reducing reagent and sample loading buffer. The mixtures were incubated at 95 °C for 5 minutes, after which they were loaded onto a Bis-Tris gel for SDS-PAGE analysis. The gel was inspected with a UV transilluminator.

### Human Embryonic Stem Cells

H9 human embryonic stem cells (WA09) were originally obtained from WiCell. H9_UBE3Am-/p-_ cells were generated and provided by Dr. Stormy Chamberlain (UCONN) [42, 43, 44]. The cells were maintained in 6-well tissue culture plates (Fisher Scientific FB012927) coated with reduced growth factor Matrigel solution (Corning 354230) at 8.7 µg/cm^2^ in mTeSR Plus medium (StemCell Technologies 100-0276). The cells were passaged using standard protocols and maintained in a humid incubator at 37°C with 5% CO_2_.

### Cell Lysate Collection

The lysates from the stem cells were collected following the manufacturers’ protocols for Pierce™ IP lysis buffer (Fisher Scientific PI87787) and M-PER™ mammalian protein extraction reagent (Fisher Scientific PI78503). Briefly, for the lysate collection with IP lysis buffer, the cells were washed with ice cold Gibco™ DPBS (Fisher Scientific 14190250). IP lysis buffer with a volume around 300-400µl was added to each well. The cells were shaken gently for 20-30min on ice. After the incubation, the wells were scraped using a cell scraper (Greiner Bio-One 541070). The cells were then centrifuged at 13,000g for 10min at 4°C. The supernatant was collected and stored in -80°C (long-term storage) or 4°C (short-term storage) until use. For M-PER reagent, briefly, the cells were washed with Gibco™ DPBS (Fisher Scientific 14190250). M-PER lysis buffer with a volume around 300-400µl was added to each well. The cells were shaken gently for 20-30min at room temperature. After the incubation, the wells were scraped using a cell scraper (Greiner Bio-One 541070). The cells were then centrifuged at 14,000g for 10min. The supernatant was collected and stored in -80°C (long-term storage) or 4°C (short-term storage) until use. Three biological replicates collected from different wells of a 6-well plate were used for Fig. 5. Amicon® ultra centrifugal filters (MilliporeSigma, UFC505096, 50kDa MWCO, 0.5ml) were used to concentrate the cell lysates. To concentrate the cell lysate, the manufacturer’s protocol was followed. The same amount of protein loss was assumed between samples prepared at the same time.

### BCA Protein Assay

Pierce™ BCA Protein Assay Kit (ThermoScientific 23225) was used to calculate the total amount of protein present in the cell lysates. The manufacturer’s protocol was followed. Briefly, the standards were prepared as suggested by the manufacturer. The BCA working reagent was prepared (50:1, Reagent A:B) based on the number samples and standards. The standards and samples with a volume of 30 µl were added to 96-well clear bottom plates (Corning, 3904). After adding 240 µL of the working reagent to each well, the plate was mixed on a shaker for 30 seconds and then incubated at 37°C for 30min in dark. The plate was then equilibrated to room temperature for 20min. The absorbance was measured at 560nm on the TECAN microplate reader. The assay was performed in technical triplicates. The outliers were removed if present. The standard curve was plotted as described by the manufacturer. A quadratic polynomial fit was used for the standard curves. The protein concentrations obtained from this assay were then used to calculate the protein amount present in the lysates. The values were adjusted depending on the volume of the samples. For the assay, the FITC readings from the lysates were equilibrated to the same protein amount after the readings were subtracted from the negative buffer control reading.

## Declaration of Interest

A technology disclosure to North Carolina State University has been filed listing LH and AJK as inventors.

## Funding Disclosure

Research reported in this publication was supported by the Foundation for Angelman Syndrome Therapeutics (FAST) under Award Numbers FT2022-008, FT2020-003, and FT2021-002.

## Acknowledgements

We thank Dr. Allyson Berent and Jennifer Panagoulias for helpful insights into the assay needs of putative therapies. We thank Dr. Stormy Chamberlain for graciously gifting the H9_UBE3A m-/p-_ cell line. We also thank Dr. Barbara Bailus for helpful technical suggestions. Fig. 1a was created with BioRender.com.

## Author Contributions

LH and AJK conceived and designed the study, drafted the article, and provided final approval. LH, AJK, and ZBY revised the article and analyzed and interpreted the data. LH and ZBY acquired the data.

## Data Availability Statement

Source data for all plots and full unadjusted gel images are in the supplemental data excel file and also publicly available at https://doi.org/10.5281/zenodo.12667127.

## Notes

### Competing Interest Statement

The authors have declared no competing interest.

