## Supplemental figures for "A High Sensitivity Assay of UBE3A Ubiquitin Ligase Activity"

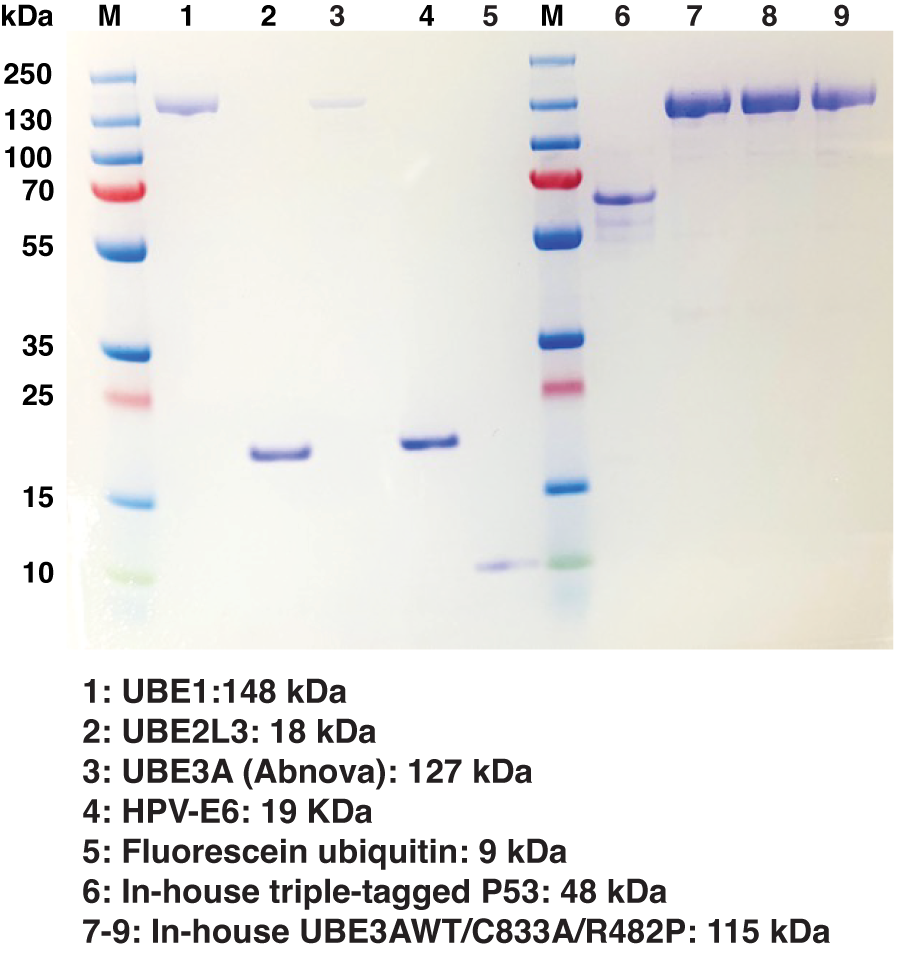

**Figure S1. SDS-PAGE of all ubiquitination assay components**

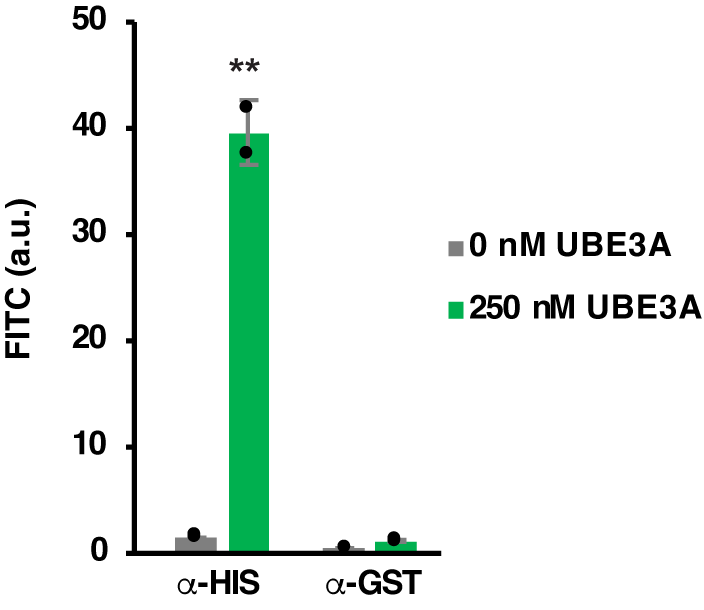

**Figure S2. UBE3A does not self-ubiquitinate to a detectable extent**

N=2 separate replicate reactions. Error bars represent standard deviation. **p<0.01 relative to sample without UBE3A. T-test with Tukey-Kramer post-hoc.

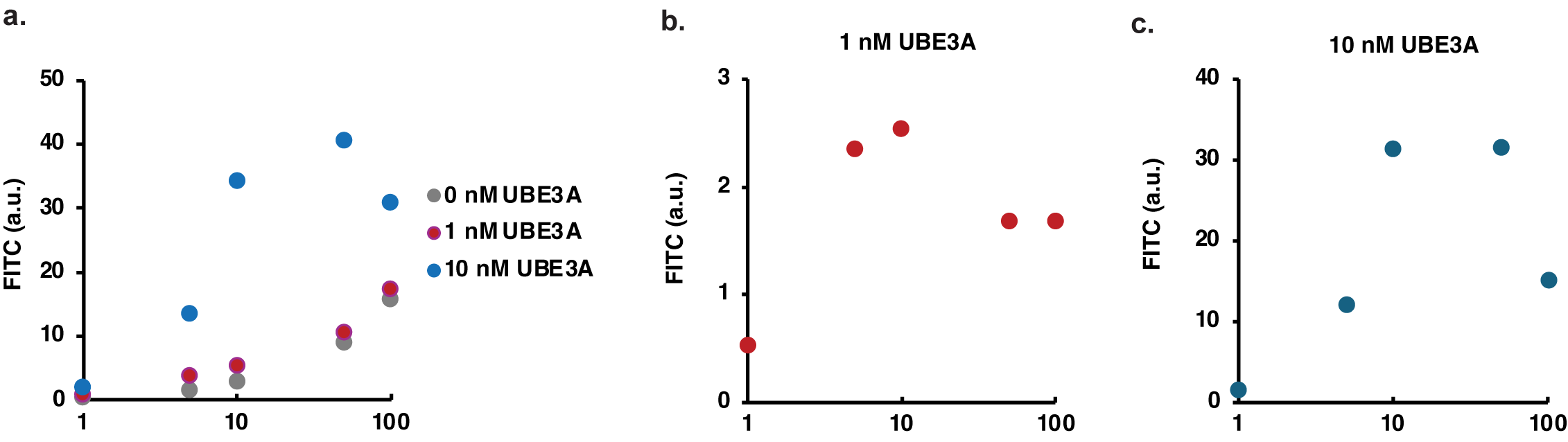

**Figure S3. Optimization of fluorescein-ubiquitin concentration**

(a) A dose-response curve was generated using 1, 5, 10, 50, 100 μM fluorescein-ubiquitin with 1 and 10 nM UBE3A separately, 0 nM UBE3A served as a negative control. (b, c) FIuorescence value of 1nM, 10nM UBE3A samples minus 0 nM (control) value.

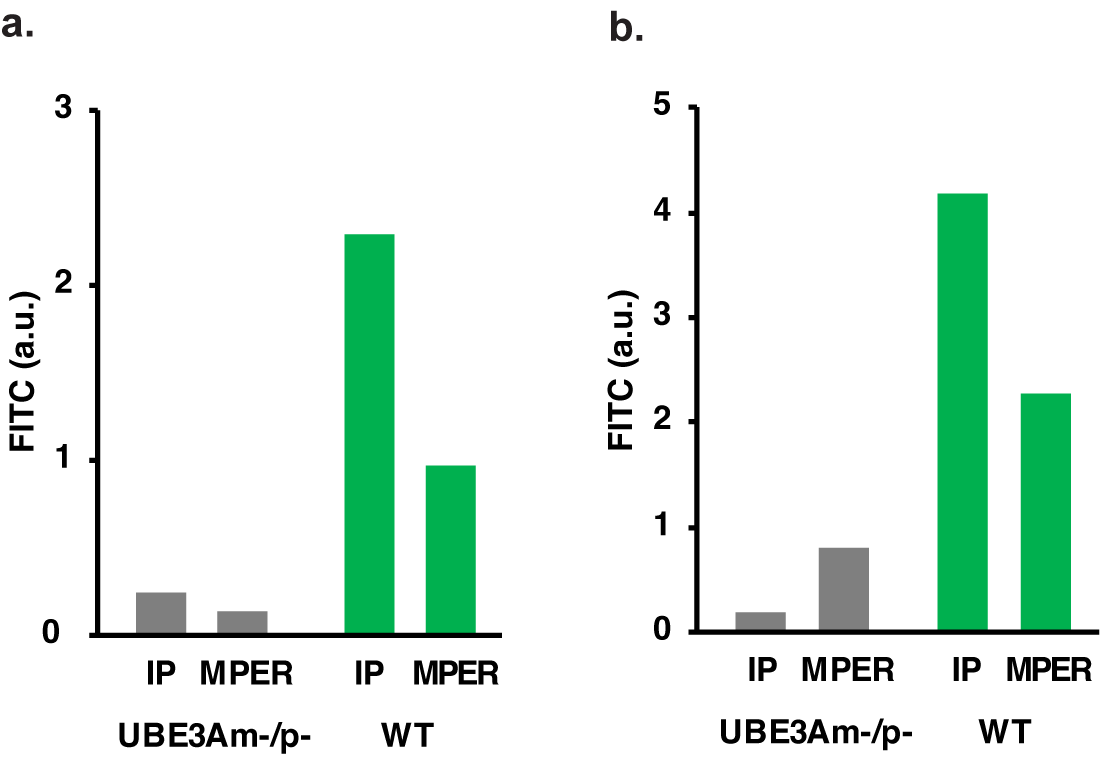

**Figure S4. Testing different lysis buffers and concentrating biological samples**

(a) Fluorescence signal from the assay performed with the cell lysates obtained from H9*_WT_* and H9*_UBE3Am-/p-_* human stem cells. IP lysis buffer and M-PER reagent were used. The fluorescence signal from the corresponding negative control (M-PER negative control or IP negative control) was first subtracted from the corresponding cell lysate signal. The fold changes between the protein amount of the corresponding sample and the lowest protein amount that a sample has (H9*_WT_* M-PER) were calculated. The signals were then equilibrated using the fold changes. Hence, all the readings were equilibrated in such a way that they all contain the same amount of protein. The negative control contains either IP lysis buffer or M-PER reagent along with other assay components except for UBE3A. N=1 biological sample collected from a well of a 6-well plate. (b) Fluorescence signal from the assay performed with the cell lysates obtained from H9*_WT_* and H9*_UBE3Am-/p-_* human stem cells. IP lysis buffer and M-PER reagent were used. The samples were filter-concentrated with ultra centrifugal filters. The FI reading from the corresponding negative control (M-PER negative control or IP negative control) was first subtracted from the corresponding cell lysate FI readings. The fold changes between the protein amount of the corresponding sample and the protein amount that the H9*_WT_* M-PER sample has were calculated. The readings were then equilibrated using these fold changes. Hence, all the readings were equilibrated in such a way that they all contain the same amount of protein. The negative control contains either IP lysis buffer or M-PER reagent (not filter concentrated) along with other assay components except for UBE3A. N=1 biological sample collected from a well of a 6-well plate.
